# Creating coveted bioluminescence colors for simultaneous myriad-color bioimaging

**DOI:** 10.1101/2024.04.28.591475

**Authors:** Mitsuru Hattori, Tetsuichi Wazawa, Mariko Orioka, Yuki Hiruta, Takeharu Nagai

## Abstract

Bioluminescence, an optical marker that does not require excitation by light, allows researchers to simultaneously observe multiple targets, each exhibiting a different color. Notably, the colors of the bioluminescent proteins must sufficiently vary to enable simultaneous detection. Here, we aimed to introduce a method that can be used to expand the color variation by tuning dual-acceptor bioluminescence resonance energy transfer (BRET). Using this approach, we could visualize multiple targets with up to 20 colors through single-shot acquisition using a color CMOS camera. Overall, this method enables simple and simultaneous observation of numerous biological targets and phenomena.

## Introduction

Labeling individual cells within a cell population is important for achieving various research objectives, such as tracking cell fate, observing cell fractionation, and identifying rare cells with distinct characteristics^1^. Optical markers, such as fluorescence and bioluminescence, are often used to distinguish not only cells in a population but also subcellular structures, cellular proteins, and physiological substances in the same sample. Although less commonly used than fluorescence, bioluminescence, which is produced by the chemical reaction of luciferin catalyzed by proteins, such as luciferase, eliminates the need for excitation light, avoids autofluorescence, and enables highly sensitive detection with a high signal-to-noise ratio^2^. Multiple luciferase color variants are essential for identifying multiple targets, and species with various bioluminescence colors have been isolated and developed^3–5^. For imaging applications, such as microscopy, the enhanced Nano-lantern^6^ (eNL), in which the luciferase, NanoLuc^7^ (Nluc), is fused with a fluorescent protein (FP) to induce bioluminescence resonance energy transfer (BRET), modifies the bioluminescence spectrum to achieve a five-color series. However, these color variants are insufficient for myriad-target observations.

When enough color variants are established, their use in bioimaging becomes the next challenge. In multiplex fluorescence imaging, the observation of more objects reduces the wavelength interval between fluorescent markers, leading to excitation crosstalk. As a result, this situation requires rigorous wavelength control and possibly computational processing, such as spectral unmixing^8–11^. Methods for distinguishing cells and organelles using six to eight different fluorescent molecules have been proposed. These methods comprise a rigorous process that involves separating the excitation and fluorescence wavelengths using a microscope and applying spectral unmixing with a software^12–14^. Although more than 15 fluorescent colors can be separated in biological specimens^11^, simultaneously observing these colors in the same sample remains limited owing to technical constraints.

The bioluminescence feature of no excitation light eliminates the concerns of excitation crosstalk in multi-labeled specimens. Thus, bioluminescence can further increase the number of colors observed in a sample. However, the relatively broader spectrum of bioluminescence compared with that of fluorescence leads to difficulty in multiplex color imaging. In addition, when multiple markers are applied, the total exposure time increases with an increase in the number of objects, and the individual objects are observed at different times. To overcome this problem, a method was employed to simultaneously detect different wavelengths by separating the optical path^15–17^. Yao et al. developed a phasor analysis method that is commonly used to distinguish spectrally similar luminophores^18^. This method enabled easy resolution of the six bioluminescent reporters in live cells via quantitative and instantaneous readouts. However, the detection and distinction of more than ten bioluminescence colors in biological observations, particularly at the cellular level, have not been reported. To emphasize the importance of observing many objects simultaneously, bioluminescence must abandon the use of switchable optical filters.

To effectively perform myriad-color observations, all wavelengths should be detected simultaneously. Recently, methods to detect bioluminescence comprise color CMOS cameras, such as smartphone cameras, to capture changes in bioluminescence color^19–21^. Similar to the capture of scenic photos with smartphones, simultaneously detecting all wavelengths enables the identification of different bioluminescence colors. Therefore, if the range of bioluminescence colors is sufficiently extensive, simultaneous observation of numerous targets can be realized.

In this study, we expanded the color palette by developing a method to modify the bioluminescence color of eNL using another fusion of FP for dual-acceptor BRET. By adjusting the BRET efficiencies between Nluc and the two FPs, we fine-tuned the bioluminescence colors to produce a series of eNL variants with 20 colors. Furthermore, although several color imaging methods generally require multiple consecutive image acquisitions while changing the optical filters, we succeeded in obtaining a time-lag-free single-shot observation of numerous colored cells using a color CMOS camera.

## Results

### Expanding the bioluminescence color hue by dual-acceptor BRET

Currently available eNLs have five color variants: CeNL (cyan), GeNL (green), YeNL (yellow), OeNL (orange), and ReNL (red) ^6^. To further increase the eNL color variants, we hypothesized that additional fusion with an FP could enhance the spectrum. Such fusion may induce another BRET that leads to a color change, not only via an increase or decrease in the original luminescence peak intensity, but also via the generation of a new peak (Fig. 1a). To demonstrate its feasibility, we fused eGFP to the C-terminus of CeNL (Fig. 1b, CeNL-eGFP). The change in the emission color of CeNL-eGFP was confirmed via a comparison with the original CeNL, GeNL, and Nluc. Upon combination with its substrate, coelenterazine-h, the bioluminescence color of CeNL-eGFP differed from that of the original based on image capture using a smartphone CMOS camera (Fig. 1c). The bioluminescence spectrum of CeNL-eGFP had an additional peak compared to that of CeNL (wavelength, 512 nm), with a decrease in the original peak (wavelength, 474 nm) (Fig. 1d). Of note, such complex spectrum is difficult to obtain by simply replacing the FP of the eNL with another. Dual-acceptor BRET might increase spectral variation in eNL.

**Figure 1.**
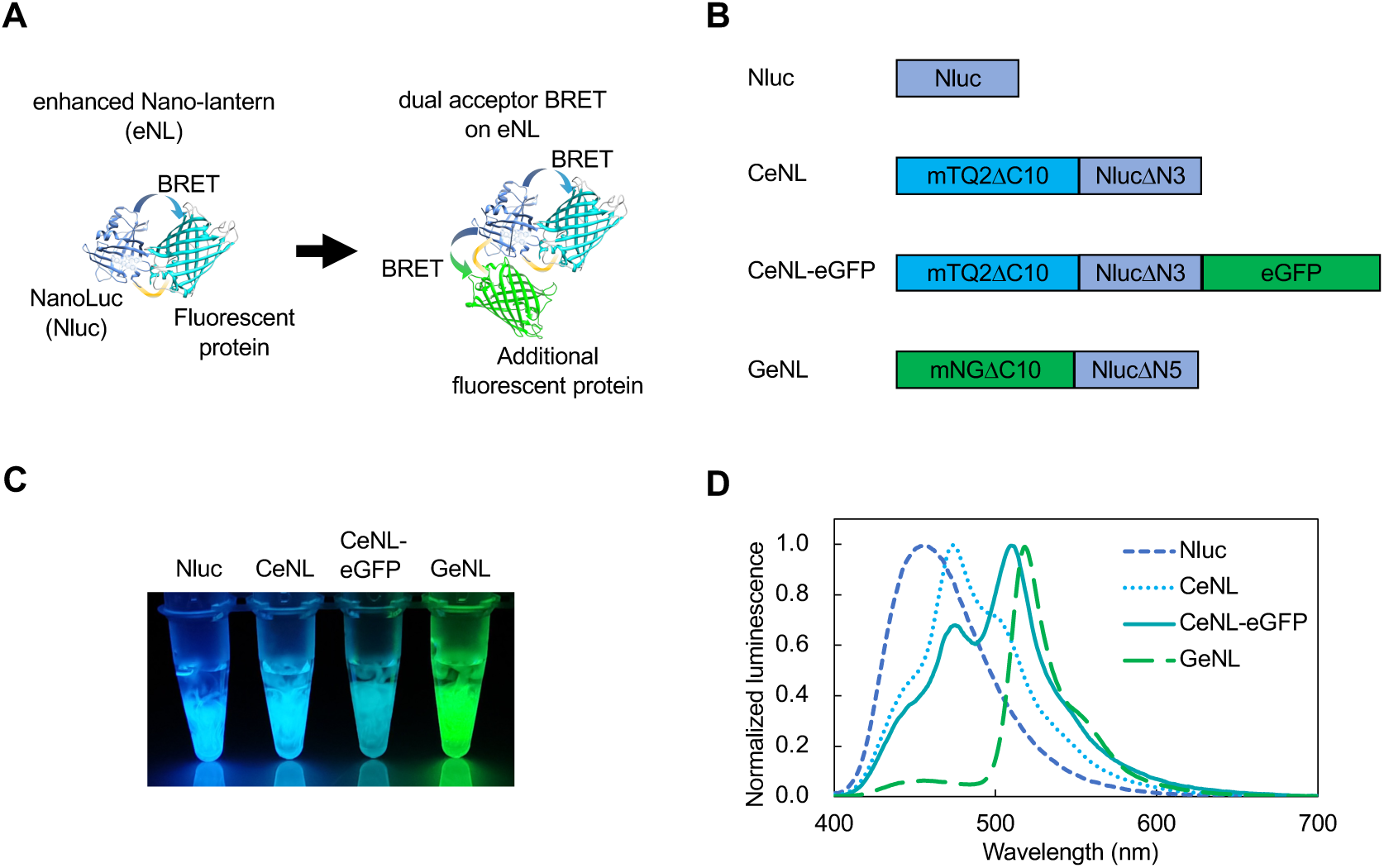
Spectrum change of eNL by the introduction of dual-acceptor BRET. **a**, Schematic of spectrum change in enhanced Nano-lantern (eNL) with the dual acceptor. **b**, Protein construction of the luciferases. mTQ2: mTurquoise2. mNG: mNeonGreen. eGFP was fused with the C-terminus of CeNL. **c**, Bioluminescence images of purified Nluc and eNL variants captured using a smartphone camera. Substrates were added to the purified proteins. **d**, Bioluminescence spectra of the eNL variants. Intensities were normalized to the peak intensity.

Based on the dual-acceptor BRET concept, we designed new eNL variants in which different types of FPs were fused to the C-terminus of the eNLs (Supplementary Fig. 1). In addition, three new eNL variants were developed based on the mCherry variants (mCXL2NL, mCXL7NL, and mCRedNL, Supplementary Note 1). When expressed in *Escherichia coli* (*E. coli*) colonies, these genes produced bioluminescence colors that were detectable by a smartphone camera (Fig. 2a). The purified protein from the *E. coli* showed 20 distinct colors (Fig. 2b), which is the largest color variation observed for bioluminescent proteins using the same luminescent substrate. The expanded color palette of the bioluminescent proteins was denoted as eNLEX (<u>eNL Ex</u>pansion). Notably, the number of colors can be further increased by changing the combination of FPs. As demonstrated by images captured with a smartphone camera, eNLEX serves as a bioluminescent marker for a myriad of targets that can be simultaneously and easily distinguished without changing the optical filters.

**Figure 2.**
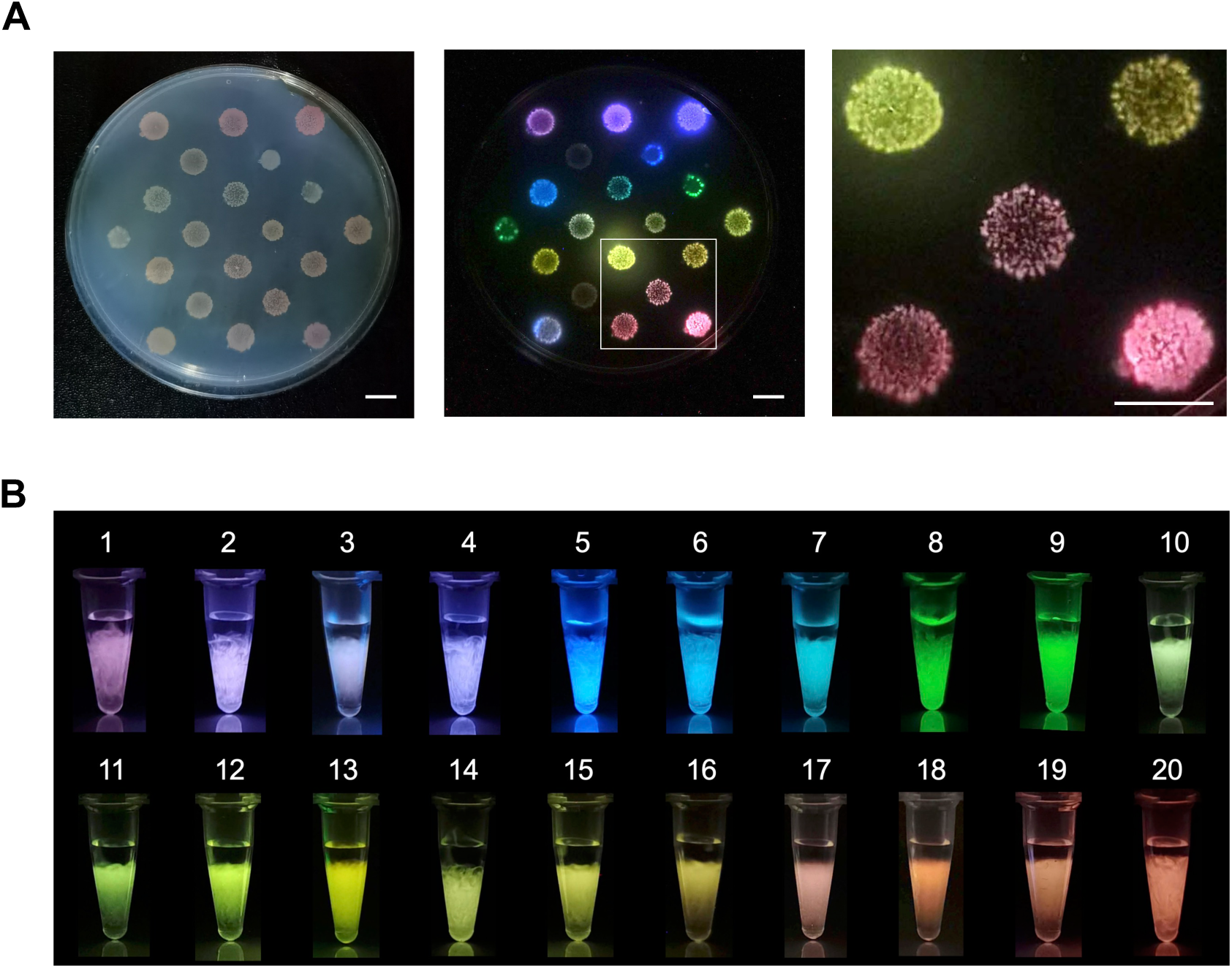
Twenty color variants of bioluminescent proteins, eNLEX. **a**, *E. coli* colonies expressing each bioluminescent protein. Brightfield image (Left), bioluminescence image (Middle), and the enlarged image of the white square (Right) are shown. Scale bar: 10 mm. **b**, Bioluminescence images of purified eNLEX members captured using a smartphone camera. 1. mCRedNL-tdT, 2. mCXL7NL, 3. CeNL-tdT, 4. mCXL2NL, 5. Nluc, 6. CeNL, 7. CeNL-eGFP, 8. GeNL, 9. YeNL, 10. OeNL-mTQ2, 11. OeNL-eGFP, 12. OeNL-Venus, 13. GeNL-tdT, 14. OeNL, 15. OeNL-mKOκ, 16. OeNL-tdT, 17. ReNL-eGFP, 18. ReNL-Venus, 19. ReNL-mKOκ, 20. ReNL.

### Slight color change of eNLEX based on adjustment of the intermolecular distance

The color variation of eNLEX was mainly due to a combination of multiple spectral peak wavelengths and differences in their peak ratios (Supplementary Fig. 2). As the intensity of each peak relies on the corresponding BRET efficiency and fluorescence quantum yield, adjusting the distance between Nluc and the FPs enables fine-tuning of subtle color differences. Using OeNL-mTurquoise2 (mTQ2) and OeNL-eGFP, we examined whether the color could be adjusted. OeNL and FP were connected via a Glu-Phe (EF) linker. To maintain a constant distance, other constructs were prepared using the rigid linker, Glu-Ala-Ala-Ala-Lys (EAAAK) (Fig. 3a). The linkers were observed to affect the bioluminescence color (Fig. 3b). For OeNL-mTQ2, the peak from mTurquoise2 (approximately 475 nm) decreased with a rigid linker (OeNL-r-mTQ2) (Fig. 3c). OeNL-eGFP exhibited three peaks derived from Nluc, eGFP, and mKOκ (Fig. 3d). Introduction of the rigid linker (OeNL-r-eGFP) induced a decrease in the eGFP peak (approximately 525 nm) and an increase in the Nluc peak (approximately 460 nm) (Fig. 3d). When the original eNLs were developed, five different colors were obtained by examining the number of deletions at each terminal of the protein and the type of amino acid used as the linker^6^. This method can be used to generate a wide range of bioluminescence colors, including minor differences, through adjustments of the length of the linker attached to the additional FP.

**Figure 3.**
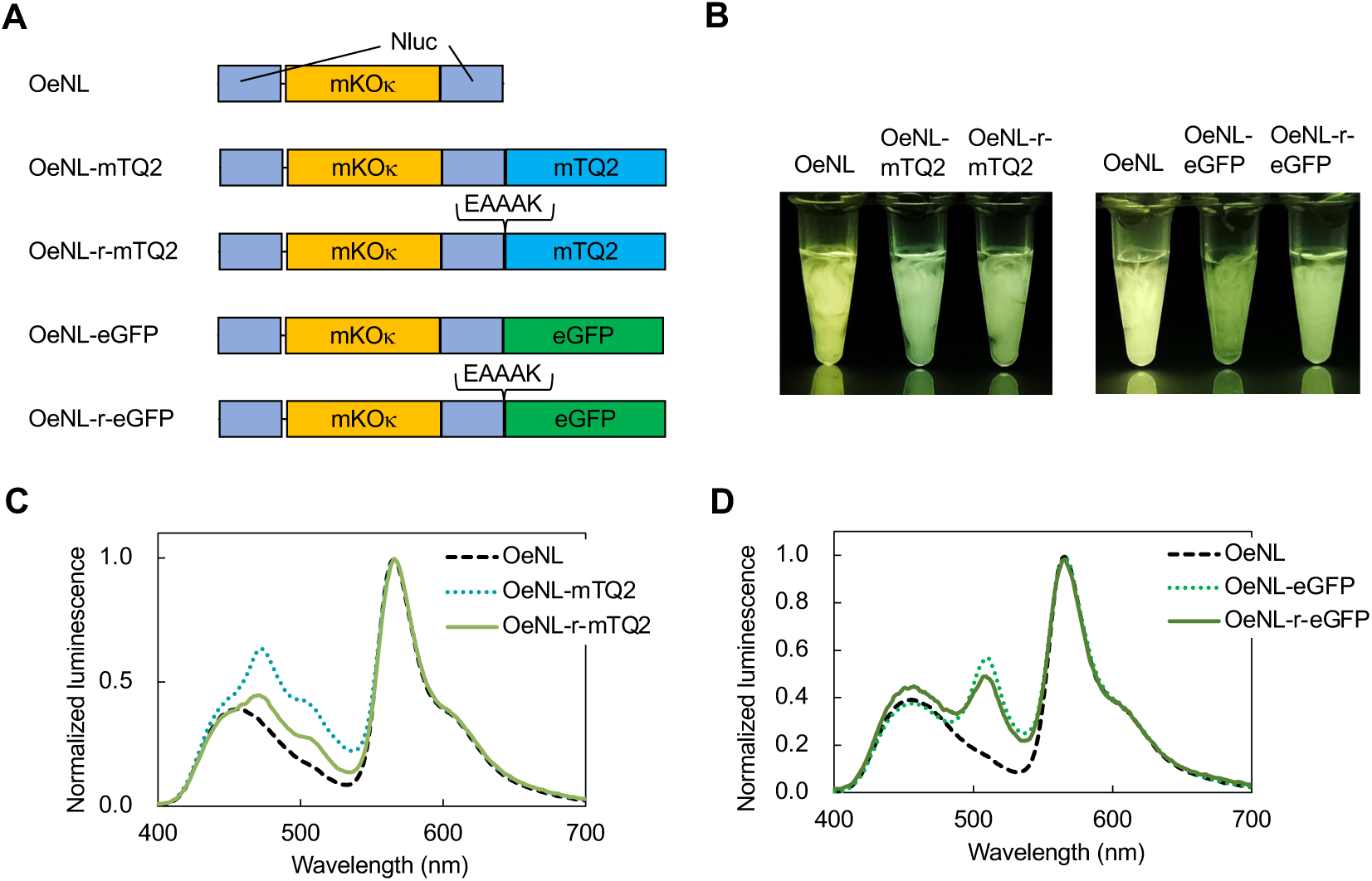
Adjusting bioluminescence colors by inserting linkers. **a**, Protein construction of OeNL, OeNL-mTQ2, OeNL-rigid(r)-mTQ2, OeNL-eGFP, and OeNL-r-eGFP. The sequence of the rigid linker is also shown. **b**, Bioluminescence images captured using a smartphone camera. **c**, Bioluminescent spectra of OeNL, OeNL-mTQ2, and OeNL-r-mTQ2. The intensities were normalized to the peak intensity. **d**, Bioluminescent spectra of OeNL, OeNL-eGFP, and OeNL-r-eGFP.

### Simultaneous myriad-color imaging of bioluminescent cells with a color CMOS camera

eNLEX can be used as a marker for multiple targets. As bioluminescence requires no excitation light, we can simultaneously detect and identify multiple wavelengths as distinct colors, as depicted in the image in Fig. 2a, which was obtained with a smartphone camera. However, owing to the insufficient intensity of bioluminescence or the limited use of highly sensitive color cameras, current microscopy methods for observing multiple bioluminescence markers typically rely on a combination of cooled EMCCD cameras and bandpass filters for sequential detection. We attempted to capture bioluminescent images of living cells using a color CMOS camera. HeLa cells expressing each of the 20 color variants of the eNLEX were prepared, and their bioluminescence was observed using a camera attached to a conventional microscope. Cells emitting 20 different bioluminescence colors were detected, despite the requirement of tens of seconds of exposure to the camera (Fig. 4a). The differences in the bioluminescence colors among the cells could be distinguished even when mixed and cultured (Fig. 4b). Importantly, the performance of bioluminescence imaging using color cameras is not limited to the eNLEX. d-Luciferin-based luciferases, such as firefly luciferase, can be captured using a color camera under certain conditions. Thus, we simultaneously captured luminescence from cells expressing Eluc^22^ and Akaluc^23^ (Supplementary Fig. 3). Based on our results, a color CMOS camera can detect multiple bioluminescence colors in a single shot, enabling enormous multiplex color imaging without a time lag.

**Figure 4.**
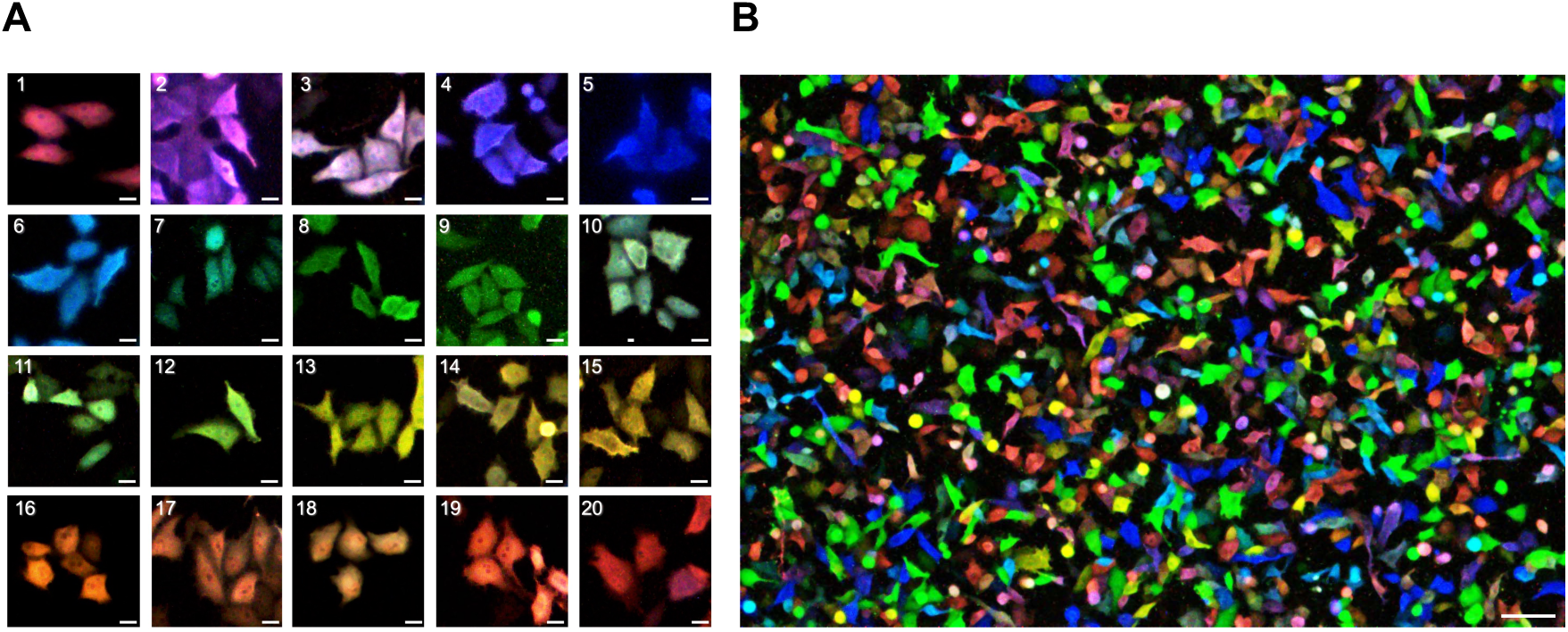
Expression of eNLEX in HeLa cells. **a**, Bioluminescence images of HeLa cells expressing each of the eNLEX members. The images were captured using a color CMOS camera. 1. mCRedNL-tdT, 2. mCXL7NL, 3. CeNL-tdT, 4. mCXL2NL, 5. Nluc, 6. CeNL, 7. CeNL-eGFP, 8. GeNL, 9. YeNL, 11. OeNL-eGFP, 10. OeNL-mTQ2, 12. OeNL-Venus, 13. GeNL-tdT, 14. OeNL, 15. OeNL-mKOκ, 16. OeNL-tdT, 17. ReNL-eGFP, 18. ReNL-Venus, 19. ReNL-mKOκ, 20. ReNL. Scale bar: 20 µm. **b**, Simultaneous myriad-color bioluminescence imaging of a cell mixture expressing each of the eNLEX members. Scale bar: 100 µm.

### Application of eNLEX to bioimaging

Simultaneous multi-color bioluminescence imaging with the eNLEX can be used to detect various biological specimens. We constructed cell spheroids to demonstrate the ability of eNLEX to detect individual cells within complex cellular structures. HeLa cells transiently expressing each of the seven color variants of eNLEX were cultured to generate spheroids. Each cell within the spheroid was observed to have a distinct bioluminescence color (Fig. 5a). Furthermore, we applied simultaneous bioluminescence imaging for cell identification and observation across various spatial scales, from complex cell structures to subcellular levels. To confirm our results, we attempted simultaneous multi-color observation of subcellular structures. Each eNLEX member was fused with a localization signal and expressed in cells. As shown in Fig. 5b and Supplementary Fig. 4, the organelles (nucleus, nucleolus, cell membrane, mitochondria, lysosome, peroxisome, and ER) were successfully visualized using different colors. At the individual animal level, bioluminescence imaging is often used instead of fluorescence imaging. However, only few reports describe the multi-color observation of bioluminescence in individual animals. To verify the observability of each eNLEX color, we subcutaneously introduced cells expressing each of the seven color variants of the eNLEX into mice. As shown in Fig 5c, we confirmed the simultaneous capture of bioluminescence signals from each eNLEX member. Thus, simultaneous bioluminescence imaging is highly applicable over a wide range of spatial scales.

**Figure 5.**
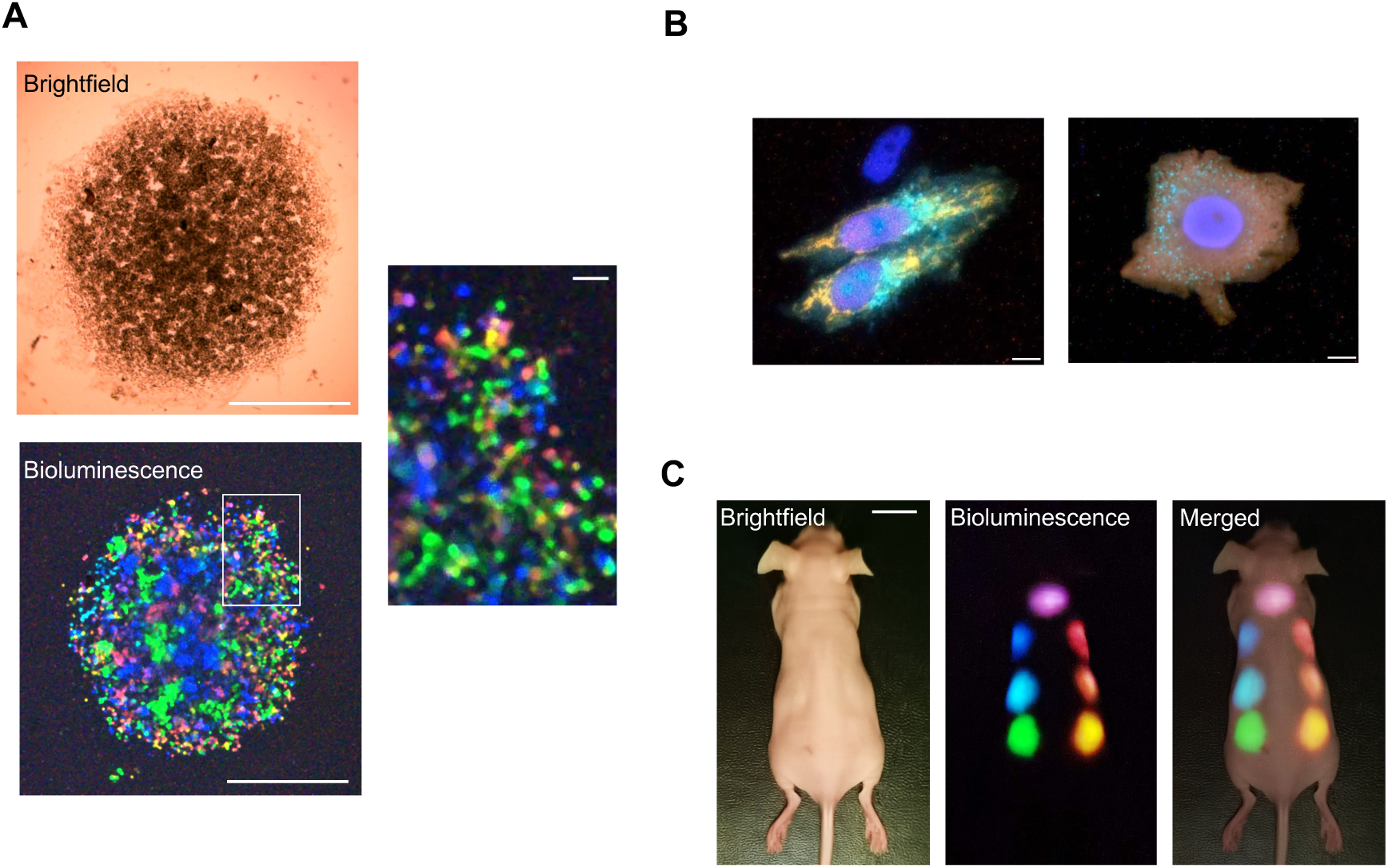
Multi-scale application of eNLEX. **a**, Brightfield (upper left) and bioluminescence imaging (lower left and right) of cell spheroid. HeLa cells expressing 7 (mCXL7NL, Nluc, CeNL-eGFP, GeNL, GeNL-tdT, OeNL-tdT, and ReNL) colors of eNLEX were cultured in the same well and observed. Scale bar: 1 mm. The enlarged image of the white square in the lower left panel is also shown in the right panel (Scale bar: 100 µm). **b**, Bioluminescence imaging of HeLa cells with eNLEX targeting the subcellular components. Left: GeNL-tdT to the mitochondria, mCXL7NL to the nucleus, CeNL to the lysosome, Nluc to the nucleolus. Right: OeNL-tdT to the cell membrane, mCXL7NL to the nucleus, CeNL to the peroxisome, ReNL to the nucleolus. Scale bar: 10 µm. **c**, Simultaneous detection of multiple bioluminescence in a mouse. Images of a mouse transfected with HEK293 cells expressing 7 (mCXL7NL, Nluc, CeNL, GeNL, GeNL-tdT, OeNL-tdT, and ReNL) color of eNLEX captured using a smartphone camera. Scale bar: 10 mm.

### Tracking the behavior of living cells through simultaneous multi-color bioluminescence imaging

When bioluminescence imaging is performed using a color CMOS camera, information regarding emission wavelengths is recorded using Red, Green, and Blue (RGB) filters on the sensor chip. This information allows the separation of each eNLEX bioluminescence color after image capture. To achieve this objective through time-course observations, HEK293 cells expressing each of the seven color variants of eNLEX were monitored for several hours. Although furimazine is an optimal luminescent substrate for Nluc^7^, its bioluminescence cannot be stably maintained over a long period owing to catalytic oxidation in the medium^24^. As a result, caged furimazines have been developed^24–27^. Here, we selected Piv-FMZ^24^ for our observations, considering the balance between the intensity and stability of bioluminescence. Using this substrate, we tracked the behavior of the cells of each color for approximately 6 h (Fig. 6a and b, Supplementary Movie 1). By converting the RGB information of the captured images into the Hue, Saturation, and Value (HSV) format^28^, cells were distinguished by their bioluminescence color (Fig. 6c, Supplementary Movie 2). Using a similar process, cells of each color in the cell spheroid image could also be distinguished (Supplementary Fig. 5).

**Figure 6.**
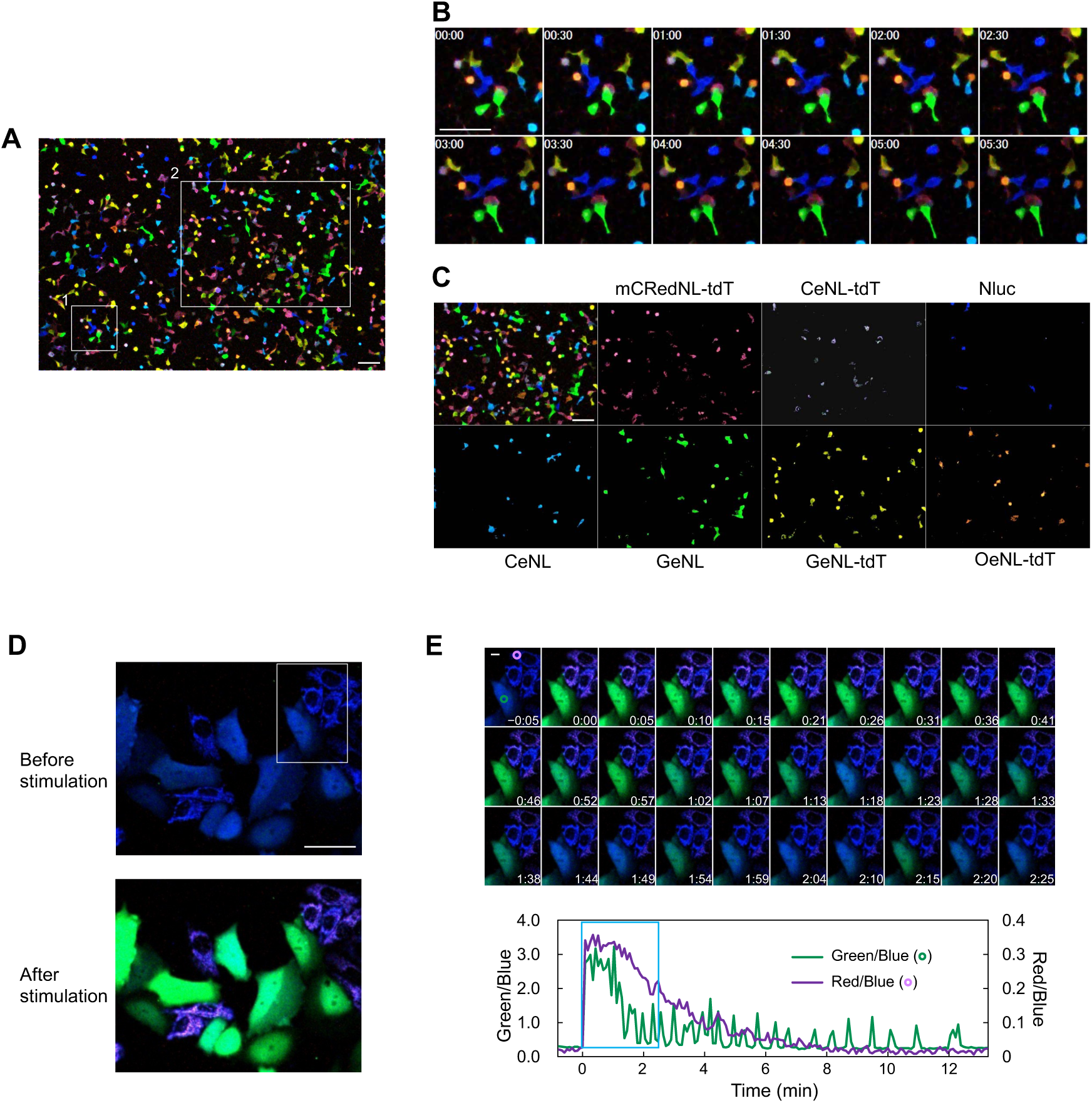
Time-lapse observation of cells via simultaneous multi-color bioluminescence detection. **a**, Simultaneous multi-color observation of HEK293 cells with bioluminescence. The movie is available as Supporting Movie 1. Each of the 7 color eNLEX members (mCRedNL-tdT, CeNL-tdT, Nluc, CeNL, GeNL, GeNL-tdT, and OeNL-tdT) was expressed in cells and cultured as a mixture. Scale bar: 100 µm. **b**, Time-course observation of cells by multiple bioluminescence color in region-1 shown in panel a. Time on images indicate the elapsed time (hours:minutes). Scale bar: 100 µm. **c**, Separation of the cell image by each bioluminescence color in region-2 shown in panel a. The emission color of each luciferase was distinguished by the RGB information. **d**, Bioluminescence imaging of intracellular Ca^2+^ changes. HeLa cells expressing bioluminescent Ca^2+^ sensors, ORCA-Y (cytosol) and ORCA-R (mitochondria), were observed using a color camera. Cells were stimulated by the addition of histamine. The white square indicates the image area in d. Scale bar: 50 µm. **e**, Time-course of Ca^2+^ based on bioluminescence detection. Time on the images indicate the elapsed time after the addition of histamine (minutes:seconds). The graph shows the luminescence changes in hue in the green and purple circle regions on the image. The blue square indicates the range of images shown. Scale bar: 10 µm.

The distinction of each bioluminescence color can be applied to monitor not only the difference between cells but also the change in the bioluminescence spectrum. Several bioluminescence indicators have been developed to detect biological targets by changing their emission color^15,16,29–31^. To demonstrate the effectiveness of simultaneous multi-color bioluminescence imaging, intracellular Ca^2+^ changes were tracked using the ratiometric Ca^2+^ bioluminescence sensors, ORCA-Y and ORCA-R^32^. In the presence of Ca^2+^, the respective bioluminescence colors changed from blue to green (ORCA-Y) and blue to purple (ORCA-R). As these bioluminescence color changes are based on BRET, they serve as good examples for testing the application of eNLEX to indicators. The sensors were found to localize in the cytosol and mitochondria. In particular, upon the addition of histamine, the bioluminescence colors in the cytosol and mitochondria changed from blue to green and blue to purple, respectively (Fig. 6d, Supplementary Movie 3). These color changes indicate transient increases and decreases in mitochondrial Ca^2+^ and cytosolic Ca^2+^ oscillations, respectively (Fig. 6e).

## Discussion

By conducting a dual-acceptor BRET, we successfully developed an eNLEX with 20 bioluminescence colors and demonstrated its application for cell identification. Altogether, eNLEX can be used to simultaneously image all 20 colors with a color CMOS camera, indicating the potential of extending this observation technique to widely available color cameras, including smartphones.

Bioluminescence, which does not require excitation light, is a potential alternative to circumvent the limitations of fluorescence imaging. However, several issues, notably the limited variety of available colors, have prevented its widespread use. Different methods can be used to change the bioluminescence spectrum of luciferase, such as the introduction of mutations in luciferase to change contact with substrates^5,23,33^ and the development of new substrates^34,35^. Using BRET combined with fluorescent molecules allows the prediction of spectral changes and the high-probability creation of variants^6, 36–40^. However, this method is limited due to the design and expansion of bioluminescence colors. Our method employs dual-acceptor BRET from Nluc using two FPs. This principle increased the variety of FPs used as acceptors and markedly expanded the number of colors that could be represented by bioluminescence. The relative intensities of multiple peaks must be controlled to express slight color differences. Here, we showed that such control can be achieved by simply changing the length or type of linkers that connect FP to Nluc. The original eNLs were developed by combining terminally truncated proteins to achieve maximum BRET^6^. Modifying these parts is also effective for further increasing the variety of bioluminescence colors. In addition to the existing eNL, we developed a “purple” eNL using a new combination of red FPs and Nluc. Further trials involving long Stokes-shifted FPs can efficiently shift the Nluc luminescence to longer wavelengths.

The low luminescence intensity, another weakness of bioluminescence compared to fluorescence, has almost been completely overcome by Nluc and its derivatives for application in microscopic imaging^6,7^. Bioluminescence imaging now offers subsecond temporal resolution. However, when multi-color imaging is performed, the emission filters must be switched to acquire each image. Therefore, multi-color imaging using the eNLEX, which has up to 20 colors, is extremely time-consuming. To overcome this limitation, we used a color CMOS camera and captured images of cells expressing 20 of the eNLEX colors through single-shot imaging.

By converting the format of the acquired images from RGB to HSV, we successfully identified cells with different bioluminescence colors. As this difference can be seen and recognized, an RGB representation of a bioluminescent image is sufficient for the human eye to identify colors. However, for automated identification, quantifying more subtle differences through the hue and saturation indices in the HSV format is effective. The combination of bioluminescence and a color CMOS camera takes a similar approach to human vision, detecting all colors simultaneously in the RGB format and later distinguishing them via information processing. Machine learning of image data allows the identification of more bioluminescence colors.

With the widespread use of smartphones, CMOS image sensors have become mainstream in camera development^41^. In fact, the use of monochrome CMOS cameras has become common for fluorescence methods used in scientific research. However, bioluminescence has not had a similar adoption. In our study, the simultaneous observation of multi-color bioluminescence was demonstrated by combining a color CMOS camera and eNLEX. The observation equipment is simpler because a switching optical filter is not required, thereby reducing cost. As the camera used in this study was not optimal for bioluminescence imaging, a more suitable color camera will be developed for future bioluminescence imaging. By improving its sensitivity and reducing the intrinsic noise, the exposure time for bioluminescence can be further shortened, potentially enabling more rapid observation. The linearity and gamma correction of RGB cameras should also be considered in quantitative measurements.

For satisfactory bioluminescence observations, a substrate and a camera must be selected. As demonstrated in this study, the use of caged substrates is effective at enabling stable observation as the oxidation caused by medium components is suppressed. If the purpose is limited to tracking rapid changes, the selection of a high-intensity substrate, such as the original furimazine, enables observations with a shorter exposure time. Thus, a desired image can be acquired using a suitable combination of camera and substrate.

Multi-color bioluminescence imaging has additional advantages over fluorescence imaging. One of the advantages of bioluminescence, which does not require excitation light, lies not only in the identification of targets but also in the absence of phototoxicity to the cells under observation. In optogenetics, which has gained popularity in recent years, light stimulation and bioluminescence work well when used within the same field of view^42,43^. For substrate addition, which has also been a bottleneck for bioluminescence observations, the introduction of an auto-luminescence system, including substrate synthesis^44,45^, into a target species is expected to be convenient. By using the method to create color variations and simultaneously observe multiple targets, the versatility of bioluminescence imaging can be further expanded.

## Methods

### Plasmid DNA

For the purification of proteins from *E. coli*, plasmids expressing NanoLuc and eNL proteins (CeNL, GeNL, YeNL, OeNL, and ReNL) were used as described elsewhere^6^. For fusion with FP at the C-terminus, cDNAs of stop codon-deleted eNL were amplified using PCR. The DNA oligonucleotides used for PCR were purchased from Hokkaido System Science. KOD-plus Neo (Toyobo Life Science) was used for PCR amplification. The amplified products were digested with *Bam*HI and *Eco*RI (TaKaRa Bio, Shiga, Japan). The fused FP genes, mTurqoise2 (mTQ2), eGFP, Venus, mKOκ, and tdTomato (tdT), were amplified by PCR and digested with *Eco*RI and *Hind*III. All samples digested with a restriction enzyme were isolated via gel electrophoresis using a FastGene Gel/PCR Extraction Kit (Nippon Genetics). Thereafter, the samples were mixed and cloned into the pRSETB vector (Invitrogen) using T4 ligase in Rapid Ligation Buffer (Promega). C-terminal-deleted mCRISPRed^46^ (mCRedDC9) and N-terminal-deleted Nluc (NlucDN5) were fused in mCRedNL-tdT, resulting in mCRedNL. C-terminal deleted mCherry-XL^47^ (mCherry-XLΔC7 or mCherry-XLΔC2) and NlucΔN5 were fused in mCXL7NL and mCXL2NL, and the constructs were named accordingly. Linkers in OeNL-r-mTQ2 and OeNL-r-eGFP were synthesized using primers. All plasmids were purified using the alkaline phosphatase method. The sequence was confirmed using dye terminator cycle sequencing with the BigDye Terminator v1.1 Cycle Sequencing kit (Thermo).

For the mammalian expression plasmids, cDNAs of eNLEX were amplified using the start codon. The fragments were then inserted into the pcDNA3 vector (Invitrogen) using *Bam*HI and *Apa*I. For transfection into mammalian cells, the plasmids were purified using the PureYield Plasmid Miniprep System (Promega). The original eNL plasmids were used in a previous study^6^ (CeNL: Addgene plasmid no. 85199, GeNL: Addgene plasmid no. 85200, YeNL: Addgene plasmid no. 85201, OeNL: Addgene plasmid no. 85202, and ReNL: Addgene plasmid no. 85203). A mammalian expression plasmid of Akaluc from RIKEN BRC (no. RDB15781) was used. A plasmid expressing Eluc was constructed based on pcDNA3, in which the original gene (Toyobo) was amplified and inserted using *Bam*HI and *Eco*RI. ORCA-Y and CoxVIIIx2-ORCA-R (a duplicated mitochondrial targeting sequence derived from the subunit-VIII precursor of human cytochrome c oxidase (Cox-VIII) at the N-terminus) were expressed by pcDNA3, which was used with *Bam*HI and *Eco*RI to insert the gene described elsewhere^32^.

eNLEX members were targeted to the mitochondria, nucleus, and cell membrane by replacing the GeNL sequence in pcDNA3-CoxVIIIx2-GeNL, pcDNA3-GeNL-H2B (a DNA-binding protein histone 2B (H2B) at the C-terminus), and pcDNA3-Lyn-GeNL (a myristoylation and palmitoylation sequence from lyn kinase at the N-terminus) described in the original paper^6^ with GeNL-tdT, mCXL7NL, and OeNL-tdT sequences, respectively. Mammalian expression vectors were used for ER, nucleolus, lysosome, and peroxisome targeting, as described previously.

### Protein expression and purification

To express the eNLEX members with an N-terminal polyhistidine tag, the *E. coli* strain JM109[DE3] transformed with the expression vector was cultured at 23 °C for 60 h in LB bacterial growth medium supplemented with 0.1 mg·mL^-1^ carbenicillin. The cultured cells were collected and disrupted by sonication (SONIFIER 150, BRANSON) after treatment with 40 µg·mL^-1^ lysozyme. The recombinant proteins were purified from the supernatants using Ni-NTA agarose affinity columns (Qiagen). The protein concentration was measured using the Bradford method (protein assay kit, Bio-Rad) and adjusted by adding 20 mM HEPES buffer (pH 7.4).

### Bioluminescent spectra measurement

The eNLEX spectra were measured using the photonic multichannel analyzer, PMA-12 (Hamamatsu Photonics), at room temperature (R.T.) with a 500 ms exposure. Coelenterazine-h (CTZh, Wako; final concentration, 5 μM) was used as the substrate.

### Detection of bioluminescence from *E. coli* using a smartphone camera

To capture the *E. coli* colonies via imaging, *E. coli* expressing each eNLEX member was colonized and covered with 2% agarose-gel containing 10 µM CTZh as droplets. Bioluminescence was detected using a camera on a smartphone (HUAWEI P40 Pro, dual-pixel sensor with 1220 × 10^4^ pixels, F-number: 1.7). Images were captured in the dark at R.T. using a preset camera software in the Android OS (manual mode, ISO250, white balance: 6200 K, exposure time: 10 s, autofocus). To capture purified eNLEX via imaging, the proteins were diluted to 1 µM with 100 µL of dH_2_O and dispended into a microtube. Twenty-five µL of 50 µM CTZh in PBS was subsequently added. The bioluminescence was captured in the dark at R.T. Images were acquired using a preset camera software (Manual mode, ISO250, white balance: 6200 K, exposure time: 0.5 s, autofocus).

### Culture and transient transfection of mammalian cells

HeLa and HEK293 cells (RIKEN BRC) were cultured in 24-well polystyrene flat-bottom dishes in Dulbecco’s modified Eagle’s medium (DMEM; Sigma) supplemented with 10% fetal bovine serum (FBS). The next day, the cells (50% confluency) were transfected with 0.5 µg·well^-1^ plasmid DNA using polyethyleneimine (PEI Max, Polysciences) and incubated for 16 h at 37 °C in 5% CO_2_. The medium was changed, and the cells were cultured for an additional 24 h.

### Preparation and bioluminescence imaging of cells

For microscopy, the transfected cells were re-cultured in collagen-coated 35 mm glass-bottom dishes for 16 h. The medium was then replaced with phenol red-free DMEM/F12 containing 10% HBSS and 1% FBS. Furimazine (Promega) was used at a 500-fold dilution for the one-shot imaging of eNLs and time-lapse imaging of Ca^2+^. For time-lapse imaging, 1 mM Piv-FMZ substrate^26^ was used at a 100-fold dilution. For Akaluc and Eluc, 1 mM Akalumine-HCl (Fujifilm-Wako) and 1 mM d-luciferin (Fujifilm-Wako) were added, respectively. For the HeLa cell spheroids, each transfected cell was cultured in a U-bottom multi-well dish for one day. The cells were collected, dispensed in the same wells, and cultured for 2 days in DMEM/F12 supplemented with 10% FBS. Furimazine was added at a 500-fold dilution immediately before observation. Bioluminescence was observed using IX-83 (Olympus) equipped with a color CMOS camera (MTR126000A-SQ, ToupTek). The condition was set using ImageView software (Bestscope) with a 7,000K white balance, 4×4 binning, and 10–50 times gain. The exposure time ranged from 15–30 s. Ca^2+^ imaging was performed for 5 s. For time-lapse cell imaging, images were captured at an exposure time of 1 min at 5-min intervals. The objective lenses (Olympus were UPlanSApo 20× (one-shot images of a single cell), UPlanFL 10× (cell population and time-lapse images), PlanApo 60× (subcellular images), and UPlanFL 4× (spheroid images). The observation condition was set at 37 °C in 5% CO_2_. For Ca^2+^ observation, cell stimulation was performed by dropping 20 µM histamine above the field of view.

The images were analyzed using the Metamorph software (Molecular Devices). The noise in the image was reduced using an average filter in the software. The color separation in Fig. 6c was performed while converting the original RGB image to Hue, Saturation, Value (HSV) using the Color Threshold program. The formula is presented in Supplementary Note 2.

### Preparation and bioluminescence imaging of mouse

Transfected HEK293 cells were detached from the culture dish and injected under the skin of BALB/c Slc-nu/nu (15–16 g, 5 weeks, female) mice. Ten µM Piv-FMZ in saline was also injected at the same site. The images were captured in the dark at R.T. using a smartphone camera and a preset camera software (Manual mode, ISO5000, white balance: 6200 K, exposure time: 10 s, autofocus).

## Acknowledgment

This study was partially supported by grants from the JST CREST Program (No. JPMJCR20H9 to T.N.), Ministry of Education, Culture, Sports, Science, and Technology (MEXT) (No. 18H05410 to T.N. and No. 21H00437 to Y.H.), and the Japan Society for the Promotion of Science (JSPS) (No. 22K05143 to M.H.).

## References

1. Ichimura, T. et al. Exploring rare cellular activity in more than one million cells by a transscale scope. Sci. Rep. 11, 16539 (2021). 10.1038/s41598-021-95930-7.

2. Massoud, T. F. & Gambhir, S. S. Molecular imaging in living subjects: seeing fundamental biological processes in a new light. Genes Dev. 17, 545–580 (2003). 10.1101/gad.1047403.

3. Viviani, V. R., Bechara, E. J. H. & Ohmiya, Y. Cloning, sequence analysis, and expression of active Phrixothrix railroad-worms luciferases: relationship between bioluminescence spectra and primary structures. Biochemistry 38, 8271–8279 (1999). 10.1021/bi9900830.

4. Viviani, V. R. et al. Cloning and molecular characterization of the cDNA for the Brazilian larval Click-beetle Pyrearinus termitilluminans luciferase. Photochem. Photobiol. 70, 254–260 (1999). 10.1562/0031-8655(1999)070<0254:camcot>2.3.co;2.

5. Loening, A. M., Wu, A. M. & Gambhir, S. S. Red-shifted Renilla reniformis luciferase variants for imaging in living subjects. Nat. Methods 4, 641–643 (2007). 10.1038/nmeth1070.

6. Suzuki, K. et al. Five colour variants of bright luminescent protein for real-time multicolour bioimaging. Nat. Commun. 7, 13718 (2016). 10.1038/ncomms13718.

7. Hall, M. P. et al. Engineered luciferase reporter from a deep sea shrimp utilizing a novel imidazopyrazinone substrate. ACS Chem. Biol. 7, 1848–1857 (2012). 10.1021/cb3002478.

8. Tsurui, H. et al. Seven-color fluorescence imaging of tissue samples based on Fourier spectroscopy and singular value decomposition. J. Histochem. Cytochem. 48, 653–662 (2000). 10.1177/002215540004800509.

9. Dickinson, M. E., Bearman, G., Tille, S., Lansford, R. & Fraser, S. E. Multi-spectral imaging and linear unmixing add a whole new dimension to laser scanning fluorescence microscopy. BioTechniques 31, 1272–1278 (2001). 10.2144/01316bt01.

10. Zimmermann, T., Rietdorf, J. & Pepperkok, R. Spectral Imaging and its applications in live cell microscopy. FEBS Lett. 546, 87–92 (2003). 10.1016/s0014-5793(03)00521-0.

11. Seo, J. et al. PICASSO allows ultra-multiplexed fluorescence imaging of spatially overlapping proteins without reference spectra measurements. Nat. Commun. 13, 2475 (2022). 10.1038/s41467-022-30168-z.

12. Valm, A. M. et al. Applying systems-level spectral imaging and analysis to reveal the organelle interactome. Nature 546, 162–167 (2017). 10.1038/nature22369.

13. Gorris, M. A. J. et al. Eight-color multiplex immunohistochemistry for simultaneous detection of multiple immune checkpoint molecules within the tumor microenvironment. J. Immunol. 200, 347–354 (2018). 10.4049/jimmunol.1701262.

14. Chen, K., Yan, R., Xiang, L. & Xu, K. Excitation spectral microscopy for highly multiplexed fluorescence imaging and quantitative biosensing. Light Sci. Appl. 10, 97 (2021). 10.1038/s41377-021-00536-3.

15. Yang, J. et al. Coupling optogenetic stimulation with NanoLuc-based luminescence (BRET) Ca++ sensing. Nat. Commun. 7, 13268 (2016). 10.1038/ncomms13268.

16. Inagaki, S. et al. Genetically encoded bioluminescent voltage indicator for multi-purpose use in wide range of bioimaging. Sci. Rep. 7, 42398 (2017). 10.1038/srep42398.

17. Morales-Curiel, L. F. et al. Volumetric imaging of fast cellular dynamics with deep learning enhanced bioluminescence microscopy. *Commun*. Biol. 5, 1330 (2022). 10.1038/s42003-022-04292-x.

18. Yao, Z. et al. Multiplexed bioluminescence microscopy via phasor analysis. Nat. Methods 19, 893–898 (2022). 10.1038/s41592-022-01529-9.

19. Arts, R. et al. Detection of antibodies in blood plasma using bioluminescent sensor proteins and a smartphone. Anal. Chem. 88, 4525–4532 (2016). 10.1021/acs.analchem.6b00534.

20. Hattori, M., Shirane, S., Matsuda, T., Nagayama, K. & Nagai, T. Smartphone-based portable bioluminescence imaging system enabling observation at various scales from whole mouse body to organelle. Sensors (Basel) 20, 7166 (2020). 10.3390/s20247166.

21. Tomimuro, K. et al. Thread-based bioluminescent sensor for detecting multiple antibodies in a single drop of whole blood. ACS Sens. 5, 1786–1794 (2020). 10.1021/acssensors.0c00564.

22. Nakajima, Y. et al. Enhanced beetle luciferase for high-resolution bioluminescence imaging. PLOS ONE 5, e10011 (2010). 10.1371/journal.pone.0010011.

23. Iwano, S. et al. Single-cell bioluminescence imaging of deep tissue in freely moving animals. Science 359, 935–939 (2018). 10.1126/science.aaq1067.

24. Mizui, Y. et al. Long-term single cell bioluminescence imaging with C-3 position protected coelenterazine analogues. Org. Biomol. Chem. 19, 579–586 (2021). 10.1039/d0ob02020f.

25. Riching, K. M. et al. Quantitative live-cell kinetic degradation and mechanistic profiling of Protac mode of action. ACS Chem. Biol. 13, 2758–2770 (2018). 10.1021/acschembio.8b00692.

26. Orioka, M. et al. A series of furimazine derivatives for sustained llve-cell bioluminescence imaging and application to the monitoring of myogenesis at the single-cell level. Bioconjug. Chem. 33, 496–504 (2022). 10.1021/acs.bioconjchem.2c00035.

27. Sakama, A., Orioka, M. & Hiruta, Y. Current advances in the development of bioluminescent probes toward spatiotemporal trans-scale imaging. Biophys. Physicobiol. 21, e21004 (2024). 10.2142/biophysico.bppb-v21.s004

28. Burger, W. & Burge, M. J. Digital Image Processing: an Algorithmic Introduction Using Java (Springer, 2008).

29. Aird, E. J., Tompkins, K. J., Ramirez, M. P. & Gordon, W. R. Enhanced molecular tension sensor based on bioluminescence resonance energy transfer (BRET). ACS Sens. 5, 34–39 (2020). 10.1021/acssensors.9b00796.

30. Itoh, Y., Hattori, M., Wazawa, T., Arai, Y. & Nagai, T. Ratiometric bioluminescent iIndicator for simple and rapid diagnosis of bilirubin. ACS Sens. 6, 889–895 (2021). 10.1021/acssensors.0c02000.

31. Hattori, M., Sugiura, N., Wazawa, T., Matsuda, T. & Nagai, T. Ratiometric bioluminescent indicator for a simple and rapid measurement of thrombin activity using a smartphone. Anal. Chem. 93, 13520–13526 (2021). 10.1021/acs.analchem.1c02396.

32. Hattori, M., Matsuda, T. & Nagai, T. Method for detecting emission spectral change of bioluminescent ratiometric indicators by a smartphone. Methods Mol. Biol. 2274, 295–304 (2021). 10.1007/978-1-0716-1258-3_25.

33. Nakatsu, T. et al. Structural basis for the spectral difference in luciferase bioluminescence. Nature 440, 372–376 (2006). 10.1038/nature04542.

34. Kuchimaru, T. et al. A luciferin analogue generating near-infrared bioluminescence achieves highly sensitive deep-tissue imaging. Nat. Commun. 7, 11856 (2016). 10.1038/ncomms11856.

35. Yeh, H. W. et al. Red-shifted luciferase-luciferin pairs for enhanced bioluminescence imaging. Nat. Methods 14, 971–974 (2017). 10.1038/nmeth.4400.

36. Saito, K. et al. Luminescent proteins for high-speed single-cell and whole-body imaging. Nat. Commun. 3, 1262 (2012). 10.1038/ncomms2248.

37. Takai, A. et al. Expanded palette of Nano-lanterns for real-time multicolor luminescence imaging. Proc. Natl Acad. Sci. U. S. A. 112, 4352–4356 (2015). 10.1073/pnas.1418468112.

38. Schaub, F. X. et al. Fluorophore-Nanoluc BRET reporters enable sensitive in vivo optical imaging and flow cytometry for monitoring tumorigenesis. Cancer Res. 75, 5023–5033 (2015). 10.1158/0008-5472.CAN-14-3538.

39. Chu, J. et al. A bright cyan-excitable orange fluorescent protein facilitates dual-emission microscopy and enhances bioluminescence imaging in vivo. Nat. Biotechnol. 34, 760–767 (2016). 10.1038/nbt.3550.

40. Goyet, E., Bouquier, N., Ollendorff, V. & Perroy, J. Fast and high resolution single-cell BRET imaging. Sci. Rep. 6, 28231 (2016). 10.1038/srep28231.

41. Vladan, B. & Oliver, S. Smartphone imaging technology and its applications. *Adv*. Opt. Technol. 10, 145–232 (2021).

42. Isomura, A., Ogushi, F., Kori, H. & Kageyama, R. Optogenetic perturbation and bioluminescence imaging to analyze cell-to-cell transfer of oscillatory information. Genes Dev. 31, 524–535 (2017). 10.1101/gad.294546.116.

43. Komatsu, N. et al. A platform of BRET-FRET hybrid biosensors for optogenetics, chemical screening, and in vivo imaging. Sci. Rep. 8, 8984 (2018). 10.1038/s41598-018-27174-x.

44. Close, D. M. et al. Autonomous bioluminescent expression of the bacterial luciferase gene cassette (lux) in a mammalian cell line. PLOS ONE 5, e12441 (2010). 10.1371/journal.pone.0012441.

45. Kotlobay, A. A. et al. Genetically encodable bioluminescent system from fungi. Proc. Natl Acad. Sci. U. S. A. 115, 12728–12732 (2018). 10.1073/pnas.1803615115.

46. Erdogan, M., Fabritius, A., Basquin, J. & Griesbeck, O. Targeted in situ protein diversification and intra-organelle validation in mammalian cells. Cell Chem. Biol. 27, 610–621.e5 (2020). 10.1016/j.chembiol.2020.02.004.

47. Mukherjee, S. et al. Directed evolution of a bright variant of mCherry: suppression of nonradiative decay by fluorescence lifetime selections. J. Phys. Chem. B 126, 4659–4668 (2022). 10.1021/acs.jpcb.2c01956.

