## Supplementary material for "Creating coveted bioluminescence colors for simultaneous myriad-color bioimaging": Supplementary Info_Hattori.pdf

Takeharu Nagai

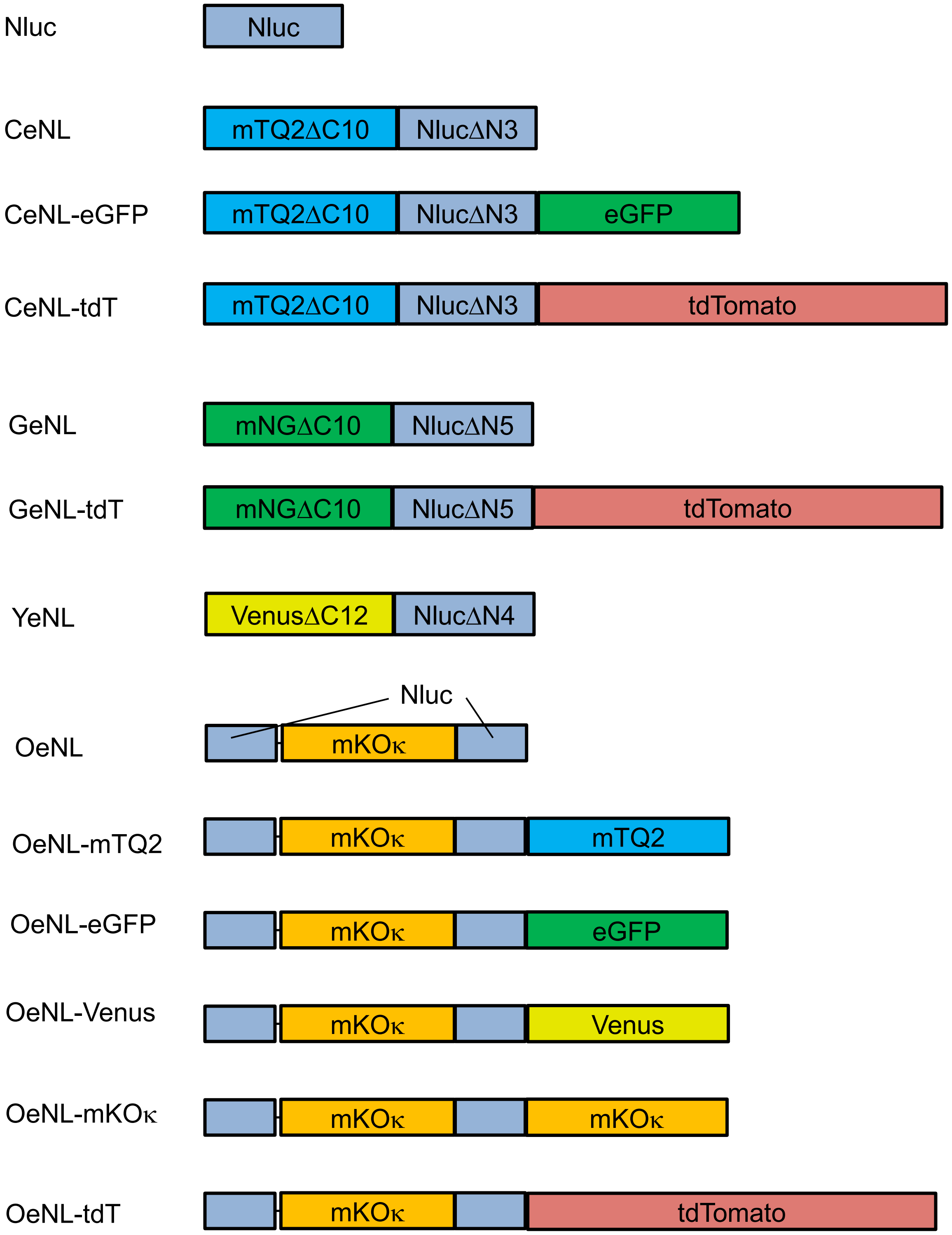
Nluc
Nluc
CeNL
mTQ2ΔC10
NlucΔN3
CeNL-eGFP
mTQ2ΔC10
NlucΔN3
eGFP
CeNL-tdT
mTQ2ΔC10
NlucΔN3
tdTomato
GeNL
mNGΔC10
NlucΔN5
GeNL-tdT
mNGΔC10
NlucΔN5
tdTomato
YeNL
VenusΔC12
NlucΔN4
OeNL
Nluc
mKOκ
OeNL-mTQ2
mKOκ
mTQ2
OeNL-eGFP
mKOκ
eGFP
OeNL-Venus
mKOκ
Venus
OeNL-mKOκ
mKOκ
mKOκ
OeNL-tdT
mKOκ
tdTomato

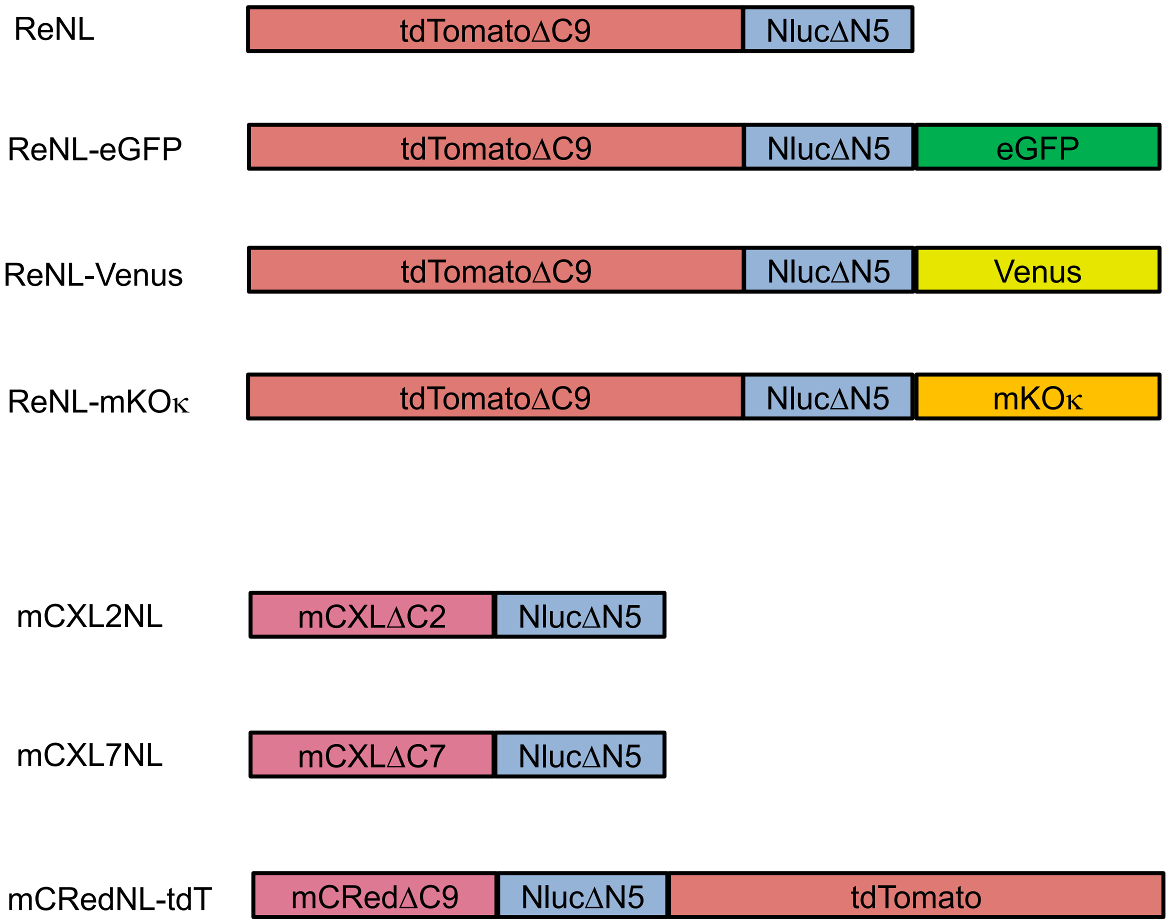


**Supplementary Figure 1. Schematic of the eNL variants.** mTQ2: mTurquoise2. mNG: mNeonGreen. tdT: tdTomato. mCXL: mCherry-XL. mCRed: mCRISPRed. Sequences from the original eNL (CeNL, GeNL, YeNL, OeNL, ReNL) were used without modification. Additional fluorescent protein was fused at the C-terminal of the original eNL by a linker of Glu-Phe (EF).

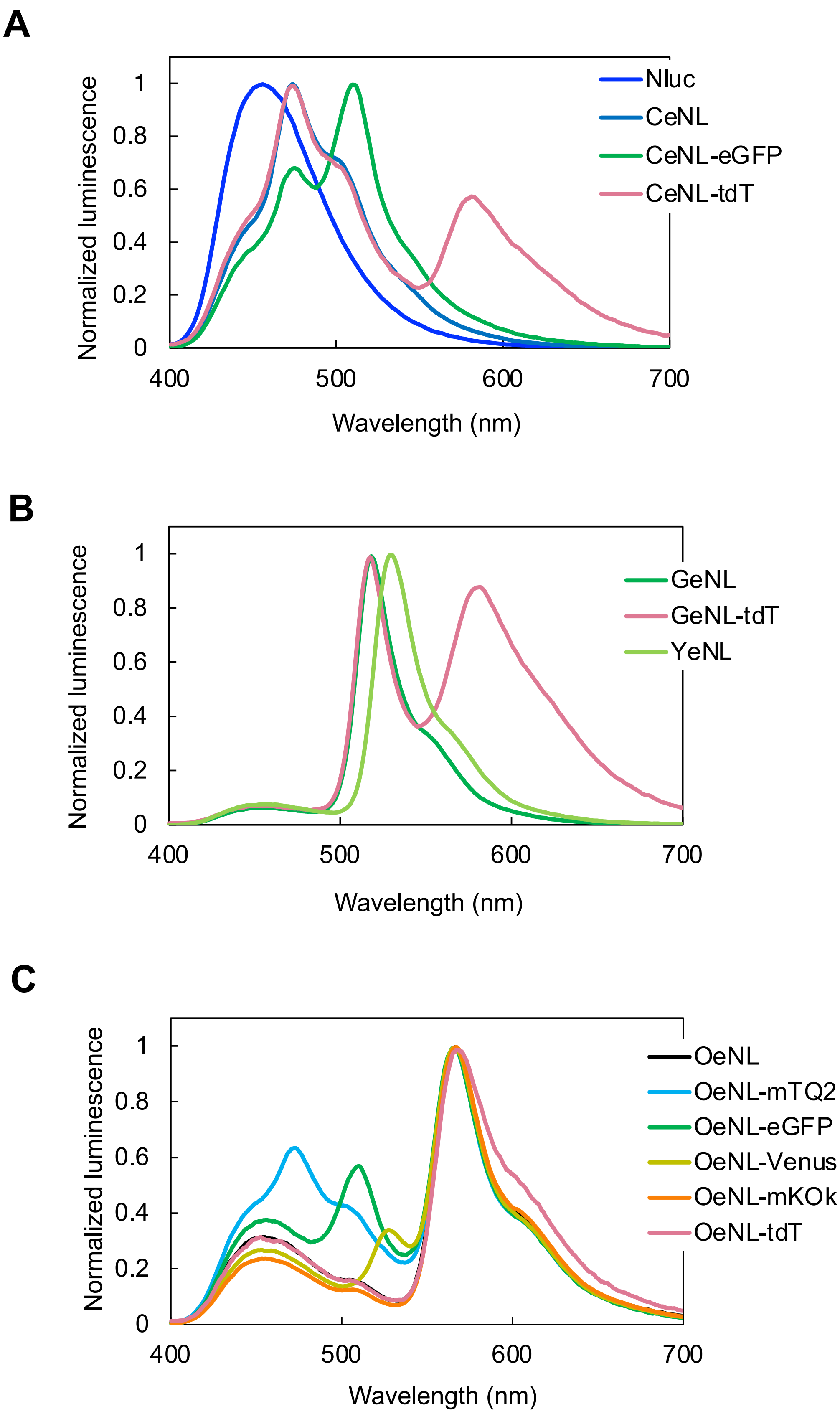

A
Normalized luminescence
Wavelength (nm)
Nluc
CeNL
CeNL-eGFP
CeNL-tdT
B
Normalized luminescence
Wavelength (nm)
GeNL
GeNL-tdT
YeNL
C
Normalized luminescence
Wavelength (nm)
OeNL
OeNL-mTQ2
OeNL-eGFP
OeNL-Venus
OeNL-mKOk
OeNL-tdT

D

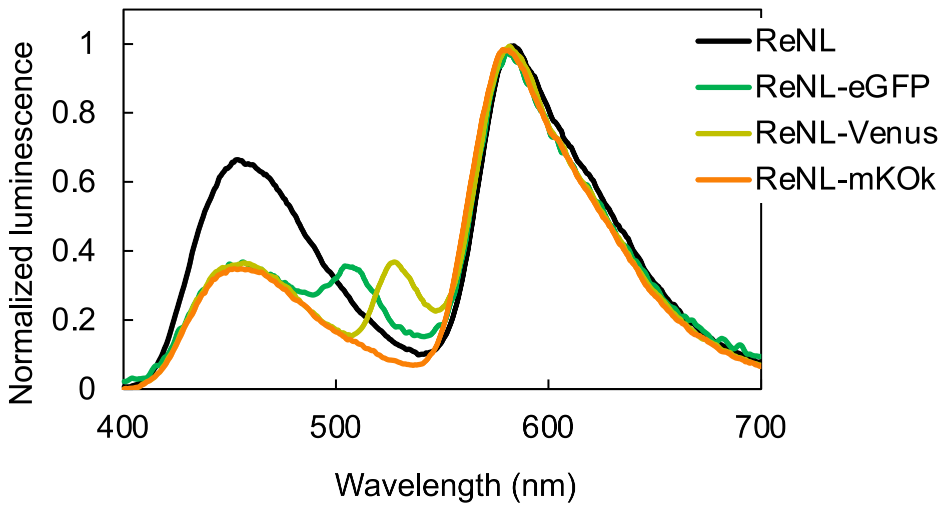


E

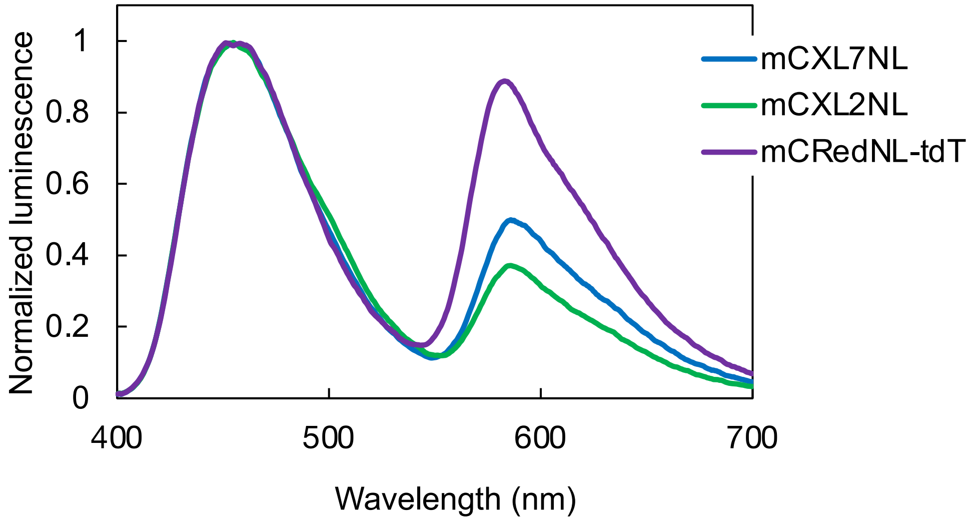


**Supplementary Figure 2. Bioluminescence spectra of eNLEX. a**, Spectra of Nluc and CeNL-based eNLs. **b**, Spectra of GeNL-based eNLs. **c**, Spectra of OeNL-based eNLs. **d**, Spectra of ReNL-based eNLs. e, Spectra of mCXL7NL, mCXL2NL, and mCRedNL-tdT. Intensities were normalized to the peak intensity.

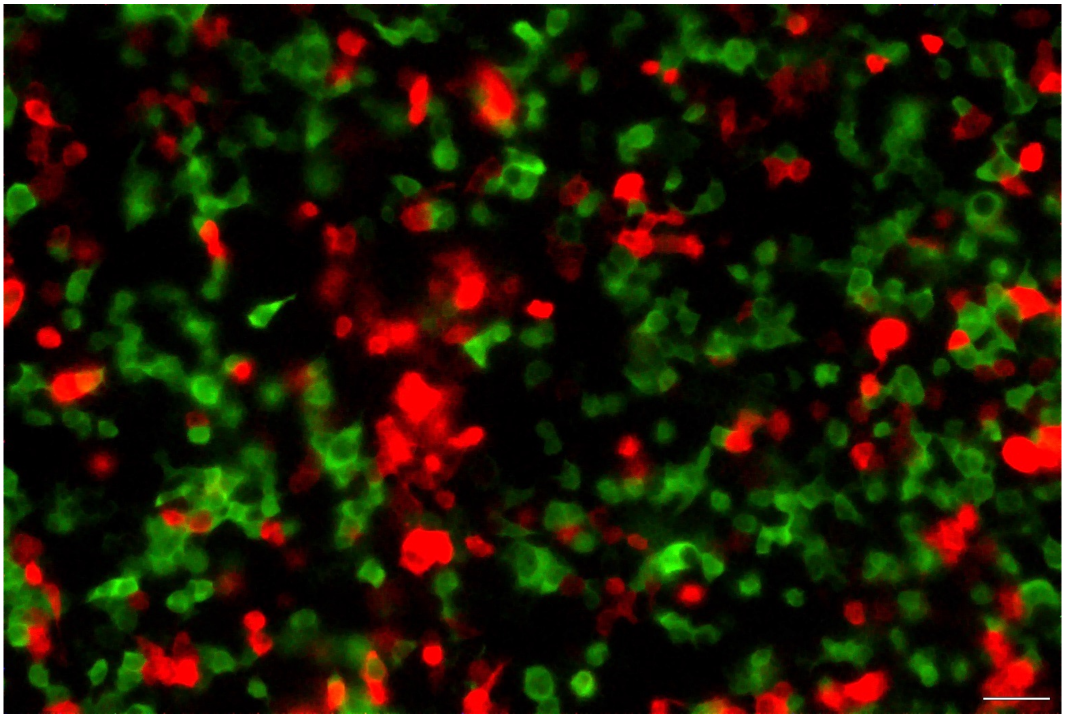

**Supplementary Figure 3. Simultaneous imaging of D-luciferin-based luciferases using a color CMOS camera.** Akalluc and Eluc were expressed in HEK293 cells. Each substrate (Akalumine-HCl and D-luciferin) was added for the emission. Exposure time: 2 min. Scale bar: 100 μm.

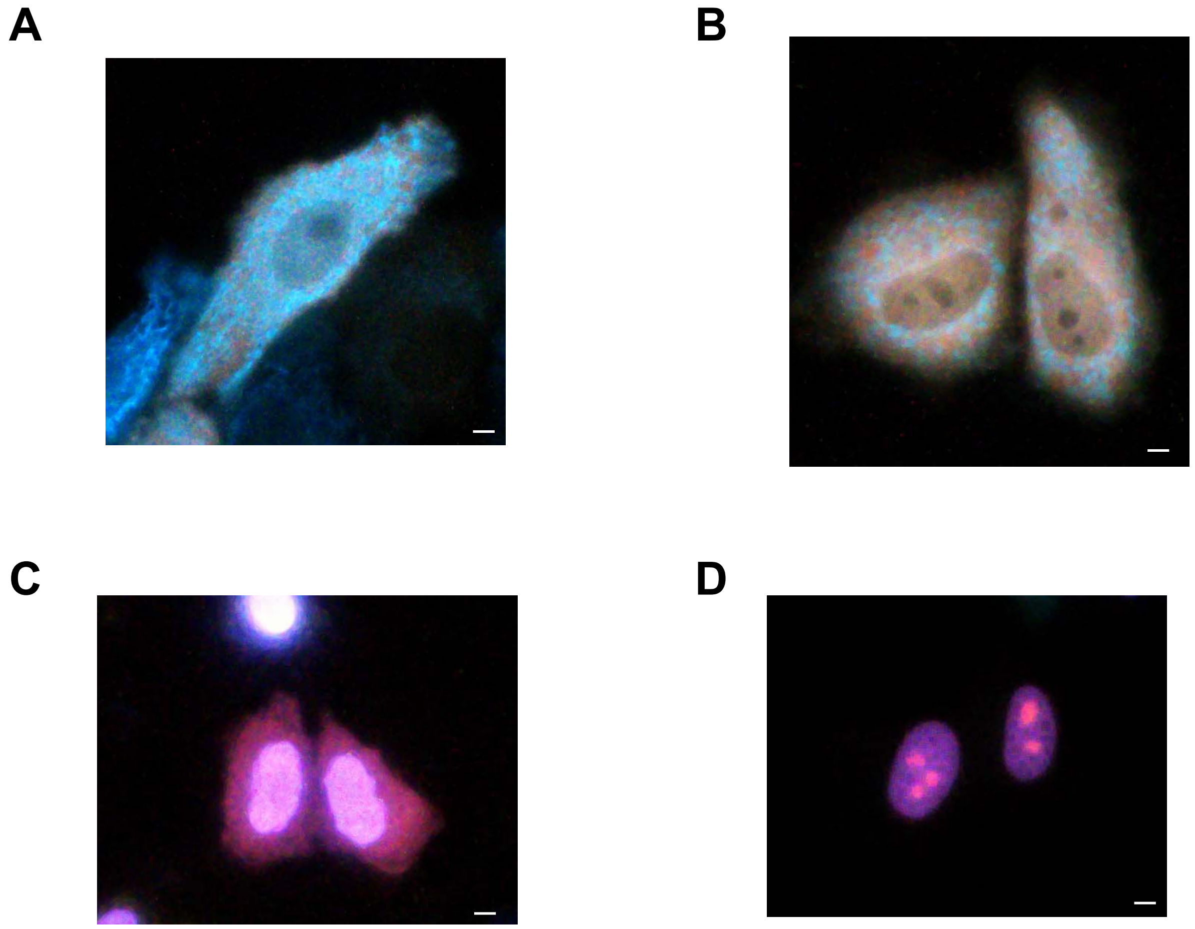


**Supplementary Figure 4. Bioluminescence imaging of HeLa cells with eNLEX targeting subcellular components. a**, CeNL to the mitochondria. OeNL-tdT to the cell membrane. **b**, Image with the same configuration but different focus. **c,** mCXL7NL to the nucleus. ReNL to the ER. **d,** mCXL7NL to the nucleus. ReNL to nucleolus. Scale bar: 10 µm.

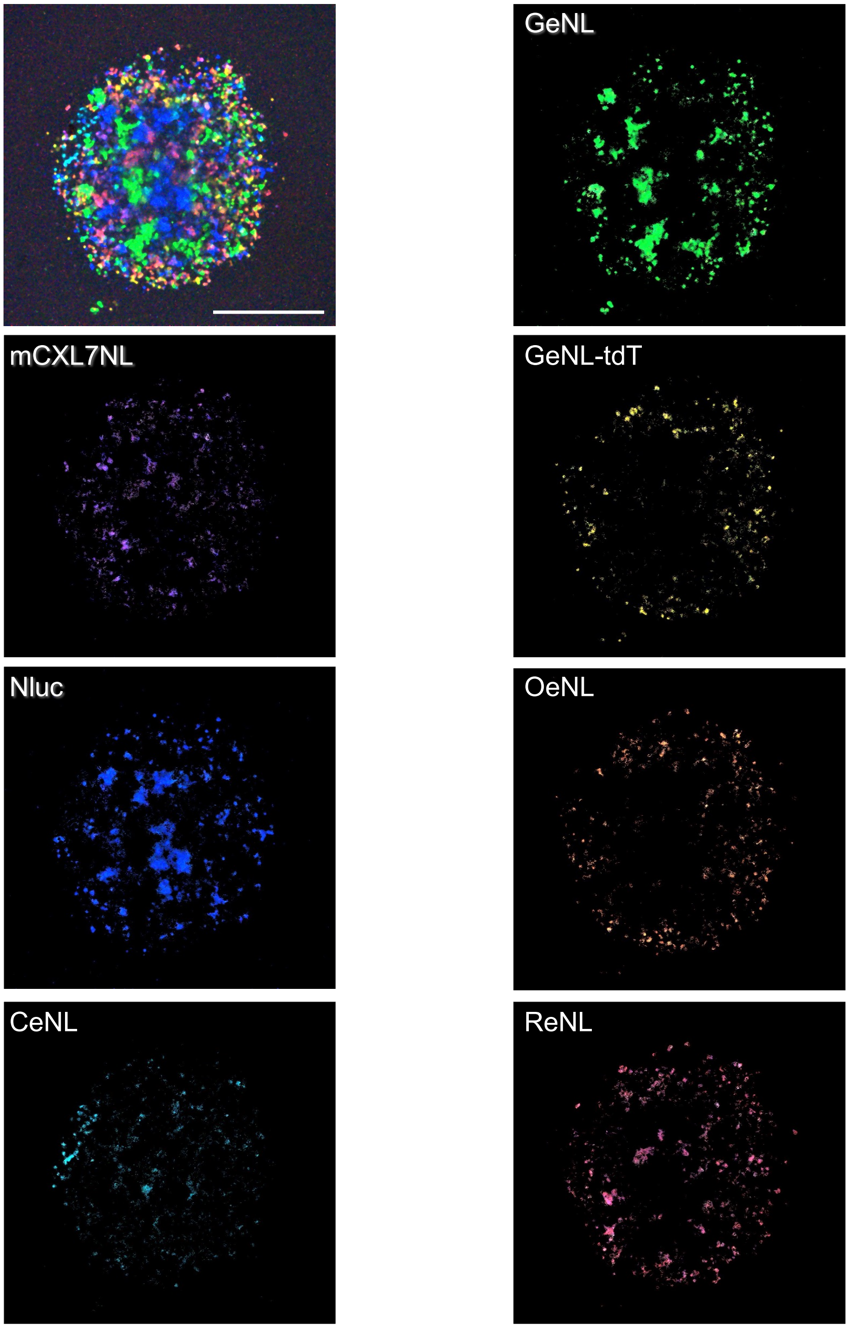


**Supplementary Figure 5. Separation of cells in the spheroid image by bioluminescence color.** The spheroid image of Fig. 5a was separated into images of each cell with respective emission colors by converting the RGB information to HSV. Scale bar: 10 mm.

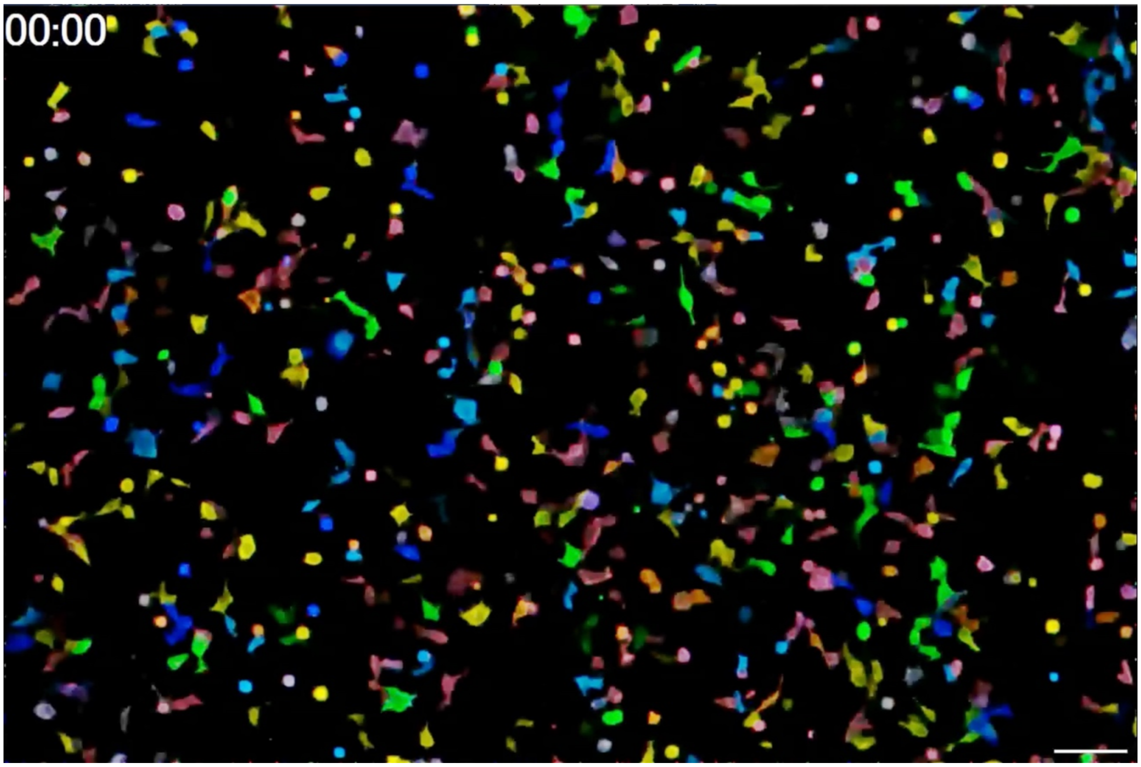


**Supplementary Movie 1. Time-course of bioluminescent cells expressing eNLEX.** Seven-colored eNLEX (mCRedNL, CeNL-tdT, Nluc, CeNL, GeNL, GeNL-tdT, and OeNL-tdT) was expressed in HEK293 cells. Images were acquired sequentially for 6 h, with 1 min of exposure. The time on the images represents the elapsed time (h: min). Scale bar, 100 μm.

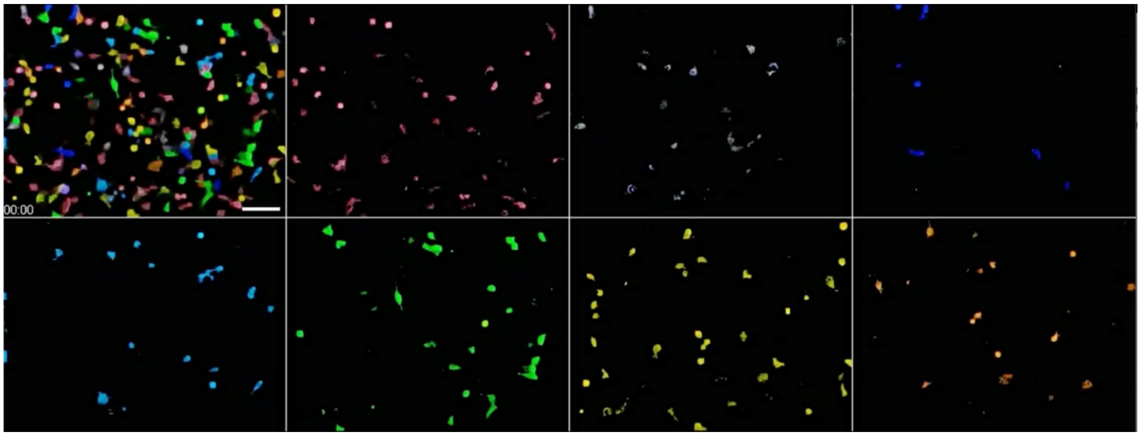


**Supplementary Movie 2. Time-course of bioluminescent cells separated by each color information.** Seven-colored eNLEX (mCRedNL, CeNL-tdT, Nluc, CeNL, GeNL, GeNL-tdT, and OeNL-tdT) was expressed in HEK293 cells. Images were acquired sequentially for 6 h, with 1 min of exposure. The time on the images represents the elapsed time (h: min). Scale bar, 100 μm.

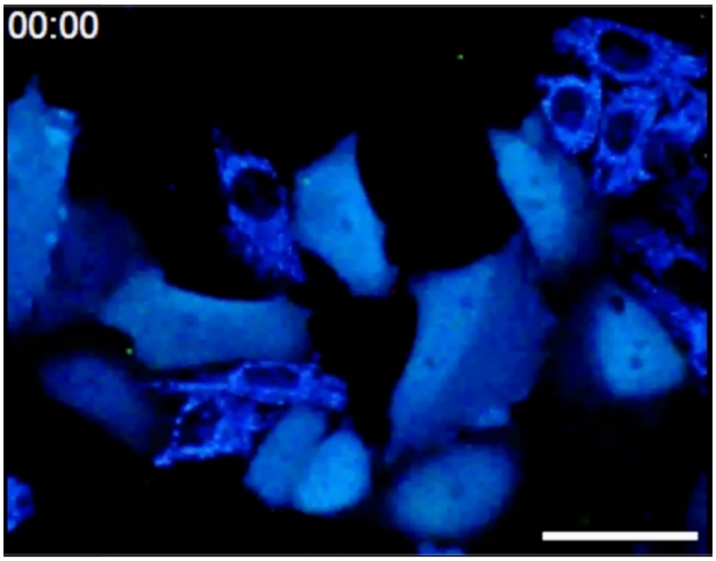


**Supplementary Movie 3. Bioluminescent $Ca^{2+}$ imaging in HEK293 cells.** ORCA-Y and ORCA-R were expressed in HeLa cells. Histamine was added 50 s after the start of the experiment. Images were acquired sequentially for 15 min, with 5 s of exposure for each image. Time on the images represents the time elapsed after the addition of histamine (minutes:seconds). Scale bar, 50 µm.

**Supplementary Note 1. Amino acid sequences of the eNL constructs newly developed in this study.** The acceptor fluorescent proteins are highlighted in yellow. Nluc is highlighted in cyan.

>mCXL2NL

MVSKGEEDNMAIIKEFMRFKVHMEGSVNGHEFEIEGEGEGRPYEGTQTAKLKVTKGGP

LPFAWDILSPQFMYGSKAYVKHPADIPDYLKLSFPEGFKWERVMNFEDGGVVTVTQDSS

LQDGEFIYKVKLRGTNFPSDGPVMQKKTMGSEASSERMYPEDGALKGEVKYRLKLKD

GGHYDAEVKTTYKAKKPVQLPGAYNVNRKLDITSHNEDYTIVEQYERAEGRHSTGGM

DELGTEDFVGDWRQTAGYNLDQVLEQGGVSSLFQNLGVSVTPIQRIVLSGENGLKIDIH

VIIPYEGLSGDQMGQIEKIFKVVYPVDDHHFKVILHYGTLVIDGVTPNMIDYFGRPYEGIA

VFDGKKITVTGTLWNGNKIIDERLINPDGSLLFRVTINGVTGWRLCERILA

>mCXL7NL

MVSKGEEDNMAIIKEFMRFKVHMEGSVNGHEFEIEGEGEGRPYEGTQTAKLKVTKGGP

LPFAWDILSPQFMYGSKAYVKHPADIPDYLKLSFPEGFKWERVMNFEDGGVVTVTQDSS

LQDGEFIYKVKLRGTNFPSDGPVMQKKTMGSEASSERMYPEDGALKGEVKYRLKLKD

GGHYDAEVKTTYKAKKPVQLPGAYNVNRKLDITSHNEDYTIVEQYERAEGRHSTGGM

GTEDFVGDWRQTAGYNLDQVLEQGGVSSLFQNLGVSVTPIQRIVLSGENGLKIDIHVIIP

YEGLSGDQMGQIEKIFKVVYPVDDHHFKVILHYGTLVIDGVTPNMIDYFGRPYEGIAVFD

GKKITVTGTLWNGNKIIDERLINPDGSLLFRVTINGVTGWRLCERILA

>mCRedNL

MVSKGEELIKENMRMKVVMEGSVNGHQFKCTGEGEGRPYEGVQTMRIKVIEGGPLPFA

FDILATSFMYGSRTFIKYPADIPDFFKQSFPEGFTWERVTRYEDGGVVTVTQDTSLEDGEL

VYNVKVRGVNFPSNGPVMQKKTKGWEADTEMMYPADGGLRGYLDRALKVDGGGHL
HCNFVTTYRSKKTVGDIKMPGVHAVDHRLERIEESDNETYVVQREVAVAKYSNLRLEDF
VGDWRQTAGYNLDQVLEQGGVSSLFQNLGVSVTPIQRIVLSGENGLKIDIHVIIPYEGLS
GDQMGQIEKIFKVVYPVDDHHFKVILHYGTLVIDGVTPNMIDYFGRPYEGIAVFDGKKIT
VTGTLWNGNKIIDERLINPDGSLLFRVTINGVTGWRLCERILA

**Supplementary Note 2. Calculation for converting RGB to HSV.**

In this study, we present a mathematical procedure for converting the RGB information into HSV. Let $R$, $G$, and $B$ be intensity readouts of the red, green, and blue channels, respectively, at a given pixel from a color camera and $0 \leq R, G, B \leq 255$, and let $H$, $S$, and $V$ be the hue, saturation, and value, respectively, at the same pixel in the HSV representation and $0 \leq H, S, V \leq 255$. MAX and MIN were defined as max($R$, $G$, $B$) and min($R$, $G$, $B$), respectively. As a result, $H$ can be expressed as follows:

$$H = (60 \times \frac{G-B}{\text{MAX}-\text{MIN}}) \times \frac{255}{360}, \text{if MAX} = R,$$

$$H = (60 \times \frac{B-R}{\text{MAX}-\text{MIN}} + 120) \times \frac{255}{360}, \text{if MAX} = G,$$

$$H = (60 \times \frac{R-G}{\text{MAX}-\text{MIN}} + 240) \times \frac{255}{360}, \text{if MAX} = B,$$

or

$$H = \text{undefined}, \text{if MAX} = \text{MIN}.$$

$S$ and $V$ were calculated using:

$$S = 255 \times \frac{\text{MAX}-\text{MIN}}{\text{MAX}},$$

and

$$V = \text{MAX}.$$
